# Homophily-informed generative models of brain maps

**DOI:** 10.64898/2026.06.07.730702

**Authors:** Vincent Bazinet, Zhen-Qi Liu, Filip Milisav, Andrea I. Luppi, Bratislav Misic

**Affiliations:** McConnell Brain Imaging Centre, Montréal Neurological Institute, McGill University, Montréal, Canada; Department of Psychiatry, University of Oxford, Oxford, United Kingdom; St John’s College, University of Cambridge, Cambridge, United Kingdom

## Abstract

The structural and functional organization of the brain can be studied across multiple scales, yielding richly detailed topographic brain maps of biological features. What are the underlying forces that shape their spatial patterning? Here we introduce a simple generative model of multiscale brain maps based on the concept of inter-regional homophily: the tendency for regions that are proximal in a given physical, molecular or functional space to display similar biological features. We evaluate the model with respect to six definitions of inter-regional homophily, including physical proximity, structural and functional connectivity, and laminar, receptor and transcriptional similarity, and across 43 empirical brain maps estimated using multiple imaging, electrophysiological and histological technologies. We show that homophilic principles are sufficient to accurately reconstruct many maps, with biological similarity and functional connectivity often contributing more than the brain’s geometry. We also identify consistent patterns of unexplained variation in maps with low homophily, revealing axes of cortical organization not captured by canonical inter-regional relationships. Finally, we show that homophily-informed generative models can be used to disentangle complex relationships between brain features and make new inferences on how they fit together. Collectively, this work highlights the fundamental contribution of homophily to the topographic layout of numerous biological features of the brain.

## INTRODUCTION

The brain is comprised of interleaved layers of organization. Transcriptomic gradients shape the arrangement of proteins, such as neurotransmitter receptors and synaptic scaffolding proteins. As the morphological properties of neurons take shape, they form nested circuits, which generate patterned electrical activity. Imaging and recording techniques increasingly allow us to take isolated snapshots of each level of organization, resulting in richly detailed brain maps. These include maps of gene transcription and cell types (sequencing) [1, 2], receptors (PET) [3, 4], microstructure (sMRI, dMRI) [5, 6], laminar differentiation (cell staining) [7, 8], metabolism (PET) [9, 10], neurophysiological rhythms (ECoG, MEG) [11–14], and haemodynamics (fMRI) [15]. Fundamentally, biological features captured by brain maps are all inter-dependent [16]. Articulating and quantifying the forces that shape the emergence of diverse brain maps is a key question in the field.

Across the cortex, brain regions that are proximal in a given space—whether physical, molecular or functional [17]—tend to exhibit similar biological properties. For example, neuronal populations that are proximal in physical space tend to be more similar to one another in terms of molecular and cellular composition, metabolism and dynamics, and ultimately, functional specialization. Inspired by the network science literature, we call this phenomenon “homophily” [18]. It reflects a general organizing principle of brain maps: features vary smoothly and coherently as a result of developmental, molecular and spatial constraints. Homophily is closely related to multiple theories of neurodevelopment, from the century-old concept of morphogen gradients [19] to the chemoaffinity hypothesis that posits that spatially patterned structures arise through local interactions of molecular and cellular signals [20]. The idea of homophily is also aligned with observations that cortical features follow continuous gradients and hierarchical axes, along which regions with similar positions share molecular, microstructural, and functional characteristics [15, 21–23]. As such, characterizing homophily across modalities offers a principled approach to quantify the shared organizational structure that underlies the interdependence of biological features.

A natural way to study homophilic tendencies in spatial maps is through the lens of autocorrelation. Indeed, the confluence of multiple physiological and geometric forces manifests at multiple levels of organization and imparts a degree of autocorrelation on virtually all maps of brain features. Traditionally, autocorrelation is thought of as similarity with respect to spatial proximity (“are brain areas that are physically close together also similar?”) but the concept is general and can be extended to other types of space (e.g. “are brain areas that are chemically close together also similar?”). The advent of multimodal imaging allows us to estimate multiple topographic cortical maps as well as multiple types of inter-regional similarity, providing a springboard from which to ask what forces shape the spatial patterning of biological brain features.

In this work, we examine how homophilic tendencies shape the distribution of biological properties across the brain. We begin by quantifying the autocorrelation of a wide array of brain maps across multiple types of inter-regional relationships, establishing a framework for understanding how geometry, connectivity and biology shape cortical organization. Building on this framework, we introduce a generative model that preserves the homophilic structure of empirical brain maps. We then use this generative model to evaluate the relative contributions of different modalities to the brain’s topographies, identify axes of cortical organization that cannot be explained by canonical inter-regional relationships, and uncover new insights into the complex relationship between brain features.

## RESULTS

We analyze 43 brain maps from the neuromaps toolbox [24], including maps of cytoarchitecture, chemoarchitecture, metabolism and oscillations (Table 1, Fig. S1). The results are organized as follows. We first evaluate the tendency for proximal brain regions to share similar properties (homophily). We evaluate this tendency with respect to six measures of inter-regional proximity, including measures of physical proximity (geodesic), connectivity (structural and functional) and biological similarity (laminar, receptor and transcriptional). We then introduce a generative model preserving the homophily structure of empirical maps and quantify the relative contribution of each modality. By looking at the residuals of our model, we extract and identify topographic patterns that cannot be explained by the known organizational forces. We finally show that the model can be used as a surrogate model, allowing us to disentangle the effects of shared organizational structure in brain map comparisons.

**TABLE 1.** Cortical brain maps included in the analysis, grouped by modality. All maps were parcellated using the Schaefer 800 atlas and rank-transformed prior to spatial analyses.

| Map name | Description | Reference |
| --- | --- | --- |
| *Density of neurotransmitter receptors and transporters (PET-derived)* |  |  |
| M1 | Muscarinic acetylcholine M1 receptor density | [67] |
| 5HT6 | Serotonin 5-HT6 receptor density | [68] |
| mGluR5 | Metabotropic glutamate receptor 5 density | [69, 70] |
| 5HT2a | Serotonin 5-HT2A receptor density | [3] |
| NAT | Noradrenaline transporter density | [71, 72] |
| CB1 | Cannabinoid CB1 receptor density | [73, 74] |
| D1 | Dopamine D1 receptor density | [75] |
| VACHT | Vesicular acetylcholine transporter density | [4, 76, 77] |
| GABAA | GABA<sub>A</sub> receptor density | [78] |
| 5HT1b | Serotonin 5-HT1B receptor density | [3] |
| H3 | Histamine H3 receptor density | [79] |
| A4B2 | Nicotinic $\alpha 4/\beta 2$ receptor density | [80] |
| DAT | Dopamine transporter density | [81, 82] |
| 5HT4 | Serotonin 5-HT4 receptor density | [3] |
| 5HTT | Serotonin transporter density | [3] |
| D2 | Dopamine D2 receptor density | [83, 84] |
| MU | μ-opioid receptor density | [85, 86] |
| KOR | κ-opioid receptor density | [87] |
| 5HT1a | Serotonin 5-HT1A receptor density | [3] |
| *Additional PET markers* |  |  |
| SV2A | SV2A density (synaptic density marker) | [88] |
| HDAC | Histone deacetylase density | [89] |
| COX1 | Cyclooxygenase-1 enzyme density | [90] |
| TSPO | Translocator protein density (neuroinflammation marker) | [91] |
| *Metabolic and vascular measures* |  |  |
| cbf | Cerebral blood flow | [9] |
| CMRO2 | Cerebral metabolic rate of oxygen consumption | [9] |
| cbv | Cerebral blood volume | [9] |
| CMRglc | Cerebral metabolic rate of glucose consumption | [9] |
| *Functional organization* |  |  |
| FC_homology | Cross-species functional homology index | [30] |
| intersubjvar | Intersubject variability in functional connectivity | [29] |
| neurosynth | PC1 of Meta-analytic functional activation maps | [92] |
| *Structural and evolutionary measures* |  |  |
| thickness | Cortical thickness | [93] |
| myelin | Intracortical myelin (e.g., T1w/T2w ratio) | [93] |
| expansion (evo) | Evolutionary cortical expansion (human vs. macaque) | [30] |
| scaling (HCP) | Cortical scaling relative to total brain size (HCP) | [94] |
| scaling (NIH) | Cortical scaling relative to total brain size (NIH) | [94] |
| scaling (PNC) | Cortical scaling relative to total brain size (PNC) | [94] |
| *Electrophysiology (MEG-derived)* |  |  |
| MEG_delta | MEG power in delta band | [14, 95] |
| MEG_theta | MEG power in theta band | [14, 95] |
| MEG_alpha | MEG power in alpha band | [14, 95] |
| MEG_beta | MEG power in beta band | [14, 95] |
| MEG_gamma1 | MEG power in low gamma band | [14, 95] |
| MEG_gamma2 | MEG power in high gamma band | [14, 95] |
| MEG_timescale | Intrinsic neural timescale (MEG-derived) | [14, 95] |

### Quantifying homophily in brain maps

We begin by comprehensively quantifying homophily across brain maps. We conceptualize homophily as the tendency for proximal brain regions to share similar biological properties. It consequently reflects the extent to which inter-regional constraints and affinities may shape the topographical organization of biological features in the brain. Fig. 1a shows a subset of brain maps ordered according to the homophily of their brain regions with respect to spatial proximity. Spatially, some maps show strong homophilic tendencies (e.g. alpha power), suggesting that strong spatial constraints shape their organization, while other maps show weaker homophilic tendencies (e.g. synapse density), suggesting that spatial constraints minimally shape their organization.

**Figure 1.**
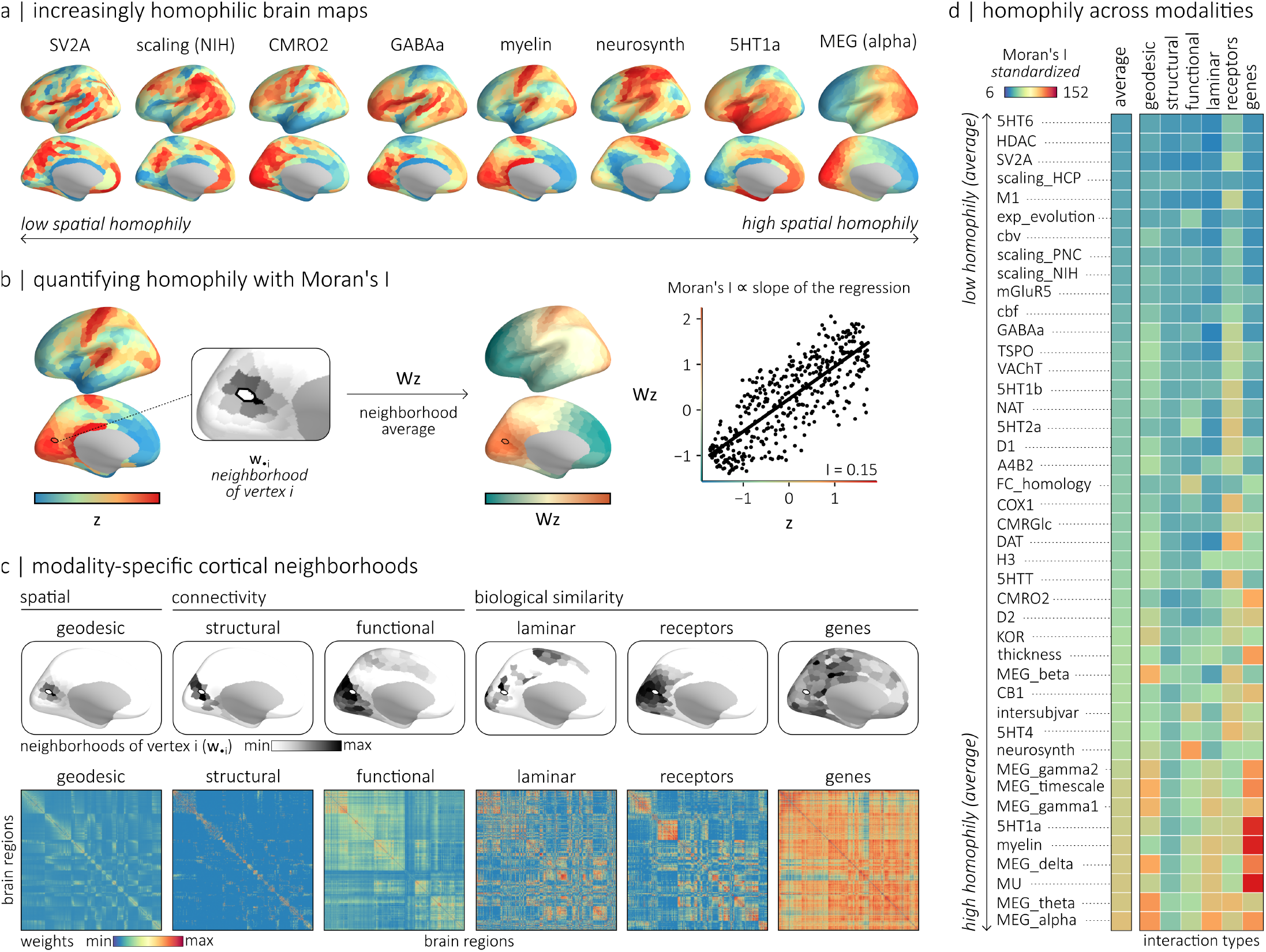
Homophily across modalities. (a) Examples of 8 brain maps, ordered according to their spatial homophily from lowest (left) to highest (right). (b) Homophily is quantified using Moran’s I. For a standardized brain map **z** and a matrix **W** capturing the relationship between brain regions, Moran’s I is estimated by relating the value in each region to the average value in its neighborhood, with the neighborhood of a region *i* given by **w**_·*i*_. More specifically, Moran’s I is proportional to the slope of the regression between the brain map **z** and the neighborhood-averaged map (**Wz**). (c) Top: examples of modality-specific neighborhoods for a region *i*. Neighborhoods can be defined by spatial proximity (geodesic), connectivity (structural or functional), or biological similarity (laminar architecture, receptor similarity or gene transcription similarity). Bottom: weight matrices encoding the strength of pairwise relationships between brain regions across modalities. (d) Homophily of 43 brain maps across inter-regional relationship types. Moran’s I values are standardized to allow comparisons across modalities (see *Methods*). Rows correspond to brain maps (ordered by average homophily), and columns correspond to relationship types. Warmer colors indicate stronger homophily, revealing systematic differences in how strongly distinct biological features align with different organizational constraints.

We operationalize homophily using a common metric for quantifying autocorrelation: Moran’s I (Fig. 1b) [25, 26]. For a standardized brain map **z**, Moran’s I is directly proportional to the slope of a regression capturing the relationship between the values in brain regions and the average values in the neighborhood of these regions (see *Methods*). Crucially, the notion of neighborhood depends on how inter-regional relationships are defined. The classical approach is to define inter-regional relationships as spatial proximity in 3D Euclidean space, but as illustrated in Fig. 1c, the same cortical region can have markedly different neighborhoods depending on whether proximity is defined geometrically, via white-matter connectivity, functional synchronization or biological similarity. In other words, cortical organization can be embedded in multiple relational spaces, each capturing distinct physiological constraints.

Fig. 1d shows the homophily of brain maps with respect to distinct inter-regional relationship. To allow comparison across modalities, Moran’s I values are standardized (see *Methods*). The cortical maps are vertically ordered according to their average homophily across modality. We find large standardized Moran’s I values with respect to transcriptional similarity for serotonergic 5HT1a (*Z*_*I*_ = 145.3) and opioid MU (*Z*_*I*_ = 152.2) receptors. This potentially explains the strong spatial co-localization between the mRNA and protein levels of these receptors, as quantified with microarray bulk sequencing, and PET imaging and autoradiography, respectively [27]. Intracortical myelin is also strongly homophilic with transcriptional similarity (*Z*_*I*_ = 148.8), which is consistent with the notion that gene expression and myelination follow a common unimodal-transmodal hierarchy [21, 28]. More generally, we find that a majority of brain maps (23 out of 43; 53%) are more strongly autocorrelated with receptor similarity than any other modality (Fig. S2). This, as expected, includes maps of neurotransmitter density, but also includes maps of inter-subject functional variability and cortical scaling. We also find that 11 maps out of 43 (26%), including maps of cortical thickness and CMRO2, are more strongly auto-correlated with transcriptional similarity than any other modality. Interestingly, only 5 maps (12%) are more strongly autocorrelated with geodesic proximity than any other modality and three of them consist of spatially smoothed MEG-derived electrophysiology maps. Altogether, these results demonstrate that the cortical distribution of a vast majority of biological features is more strongly aligned with the transcriptional and chemoarchitectural layout of the cortex than with its spatial axes.

### Modeling the autocorrelation structure of brain maps

Homophily is thought to drive the emergence of multiple biological features. In the previous section we showed that multimodal, multiscale brain maps show rich patterns of homophily with respect to various interregional relationships. Here we ask whether interregional homophilic principles are sufficient to model empirically realistic brain maps. For a given empirical map (**x**), we start with a vector of random values (**y**^(0)^)(Fig. 2a, left). Using simulated annealing, we randomly permute values while minimizing the difference in Moran’s I (Δ*I*) between the empirical and simulated map, for all six inter-regional relationships simultaneously. Fig. 2a (right) shows an example output of this generative model, with the empirical and surrogate maps side by side.

**Figure 2.**
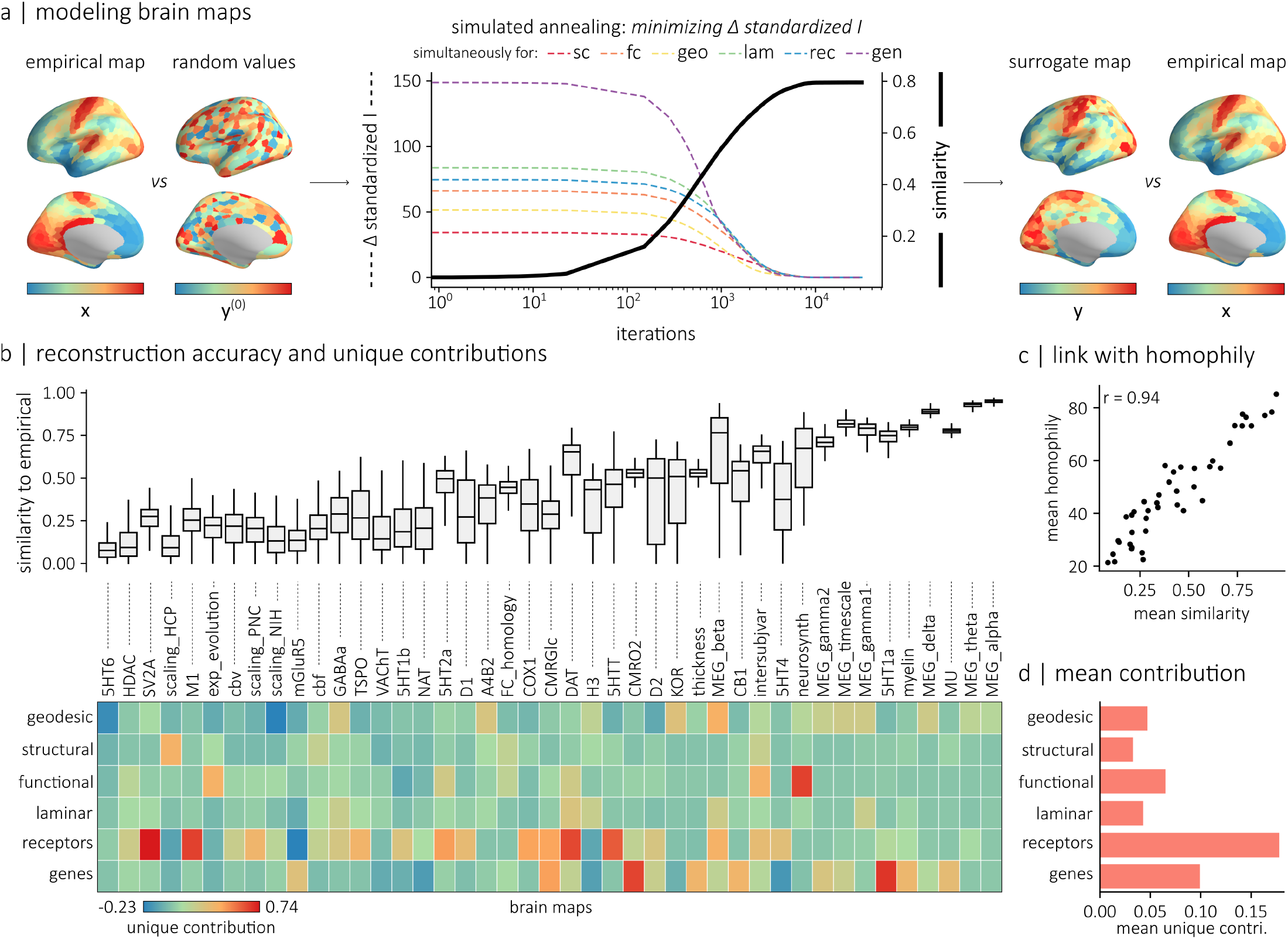
Modeling the autocorrelation structure of brain maps. (a) We introduce a generative model that constructs simulated brain maps (**y**) that preserve the homophilic structure of a given empirical map **x**. The model is initiated with a vector of random values **y**^(0)^. Pairs of values are then randomly permuted according to a simulated annealing procedure relying on an energy function that aims at minimizing the difference in the Moran’s I (Δ*I*) between simulated and empirical maps for multiple types of inter-regional relationships simultaneously. Ultimately, our model generates a simulated brain map **y** with the same homophilic structure as the empirical map. (b) Top: similarity (absolute correlation) between empirical maps and 500 simulated surrogates. Maps are ordered according to their average homophily. Bottom: relative contribution of each modality, estimated using a leave-one-out analysis. (c) Relationship between reconstruction accuracy and the mean homophily (averaged Moran’s I across modalities) of the empirical maps. (d) Average contribution of each type of inter-regional relationship across all empirical brain maps.

We generate 500 surrogate maps for each of the 43 empirical maps and evaluate the fidelity of the generative model by computing the absolute correlation between the empirical and simulated maps (Fig. 2b, top). Importantly, the average homophily of a brain map across the six inter-regional relationships is strongly correlated with the accuracy of the simulations (*r* = 0.94; Fig. 2c). This is expected, as greater homophily imposes stronger constraints on the topography of the brain map.

To explore the importance of each modality in shaping brain map organization, we performed a leave-one-out analysis in which each weight matrix was removed from the model in turn and model performance was reevaluated. The unique contribution of a modality was then defined as the relative decrease in performance compared with the full model (Fig. 2b, bottom). Fig. 2d shows the average unique contribution of each modality across brain maps. On average, the largest contributions arise from receptor similarity (mean *C*_unique_ = 0.178), transcriptional similarity (mean *C*_unique_ = 0.099) and functional connectivity (mean *C*_unique_ = 0.065). We also compared the relative model performance when using each matrix individually (i.e. their marginal contribution; Fig. S3a, b) to the unique contribution of each matrix, obtaining a metric quantifying the redundant contribution of each modality (Fig. S3a, b). A large redundant contribution indicates that, although a modality can individually constrain brain map organization, much of its explanatory power is shared with other modalities in the full model. On average, across all cortical maps, we find that geodesic weights show the largest redundant contribution (Fig. S3c). In other words, when connectivity and biological similarity is explicitly incorporated into the model, the unique contribution of geodesic weights is markedly reduced, suggesting that the predictive power of the brain’s geometric embedding mostly reflects underlying biological constraints. Biological modalities do provide unique information, indicating that they them-selves capture aspects of cortical organization that cannot be accounted for by geometry alone.

To visualize how the modeling procedure samples the wider space of possible maps and captures the unique organizational features of each map, we embed the surrogate maps in a morphospace spanned by the first two principal components of the empirical maps (Fig. 3a). We highlight 8 distinct clusters of surrogate maps, each associated with a specific empirical map. Surrogates tend to cluster near the position occupied by their representative empirical maps, with the pairwise distance between surrogate maps in this morphospace recapitulating the pairwise distance between empirical maps (*r* = 0.84, Fig. 3b). This analysis confirms that the generative model samples the unique and biologically realistic space associated with each map.

**Figure 3.**
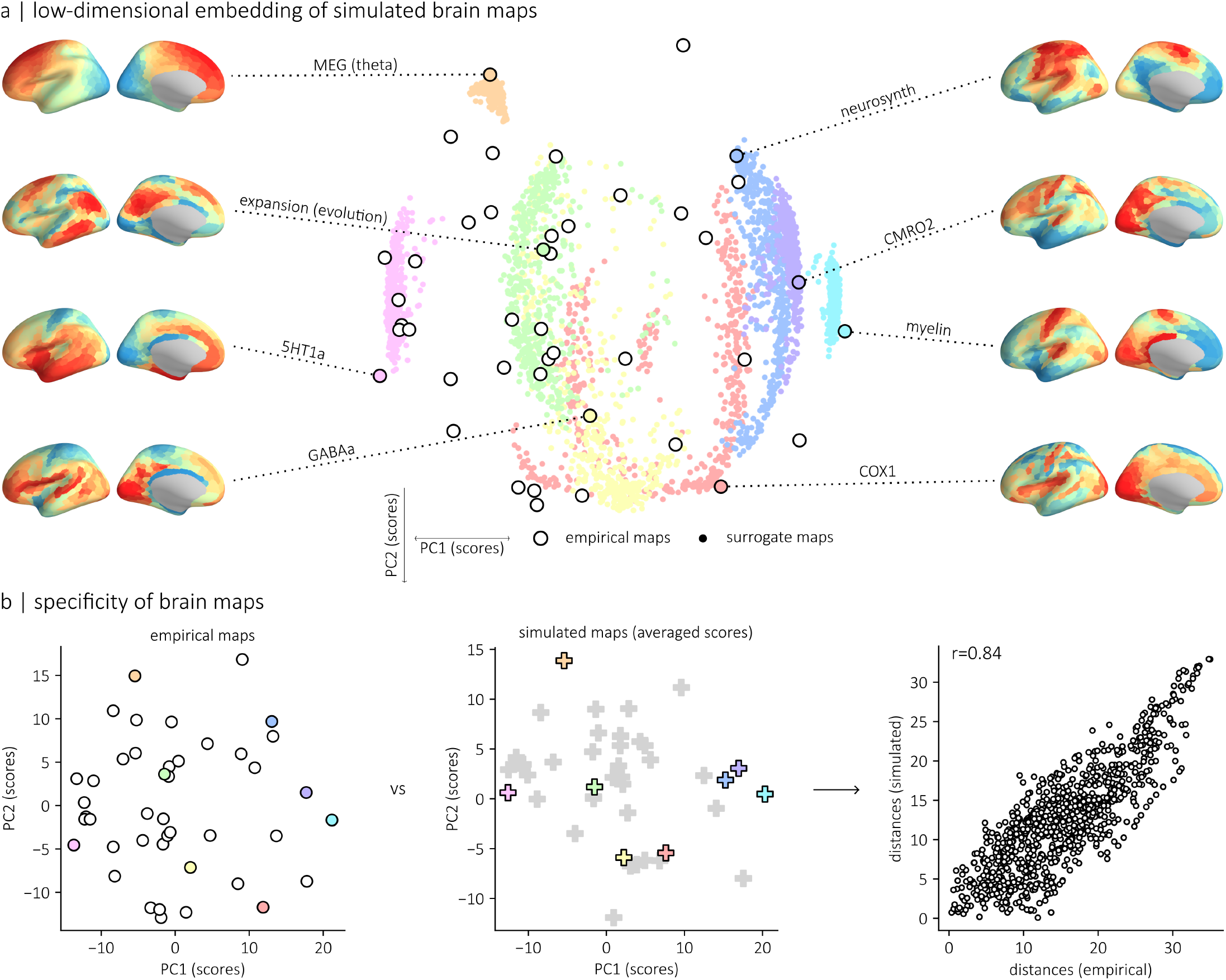
Morphospace of empirical and simulated brain maps. (a) Low-dimensional embedding (morphospace) of empirical and simulated brain maps obtained by projecting them onto the first two principal components (PC1 and PC2) of the 43 empirical brain maps used in this study. Outlined circles denote empirical maps. Clusters of simulated maps (colored points) associated with 8 distinct empirical maps are depicted. (b) The pairwise distances between the scores of empirical maps in this morphospace (left) were related to the pairwise distances between the average scores of the simulated maps associated with each empirical map (middle), revealing a strong correspondence (*r* = 0.84; right).

### Identifying patterns of unexplained variation in brain maps

In the previous section, we showed that generating brain maps with homophilic constraints recapitulates the topographical organization of empirical data. However, maps with low homophily across the six inter-regional interaction matrices cannot be accurately reconstructed.

Indeed, the organization of these maps is not constrained by either spatial proximity, structural and functional connectivity, or transcriptional, neurochemical and laminar similarity. A key question is therefore: what are the topographical patterns that cannot be explained by homophilic constraints from the modalities listed above? To answer this question, we quantified the regional errors for each empirical map as the difference between the ranked empirical values and the ranked values of the average of the simulated maps. Fig. 4a shows the regional errors for each brain map, with maps ordered according to their overall homophily.

**Figure 4.**
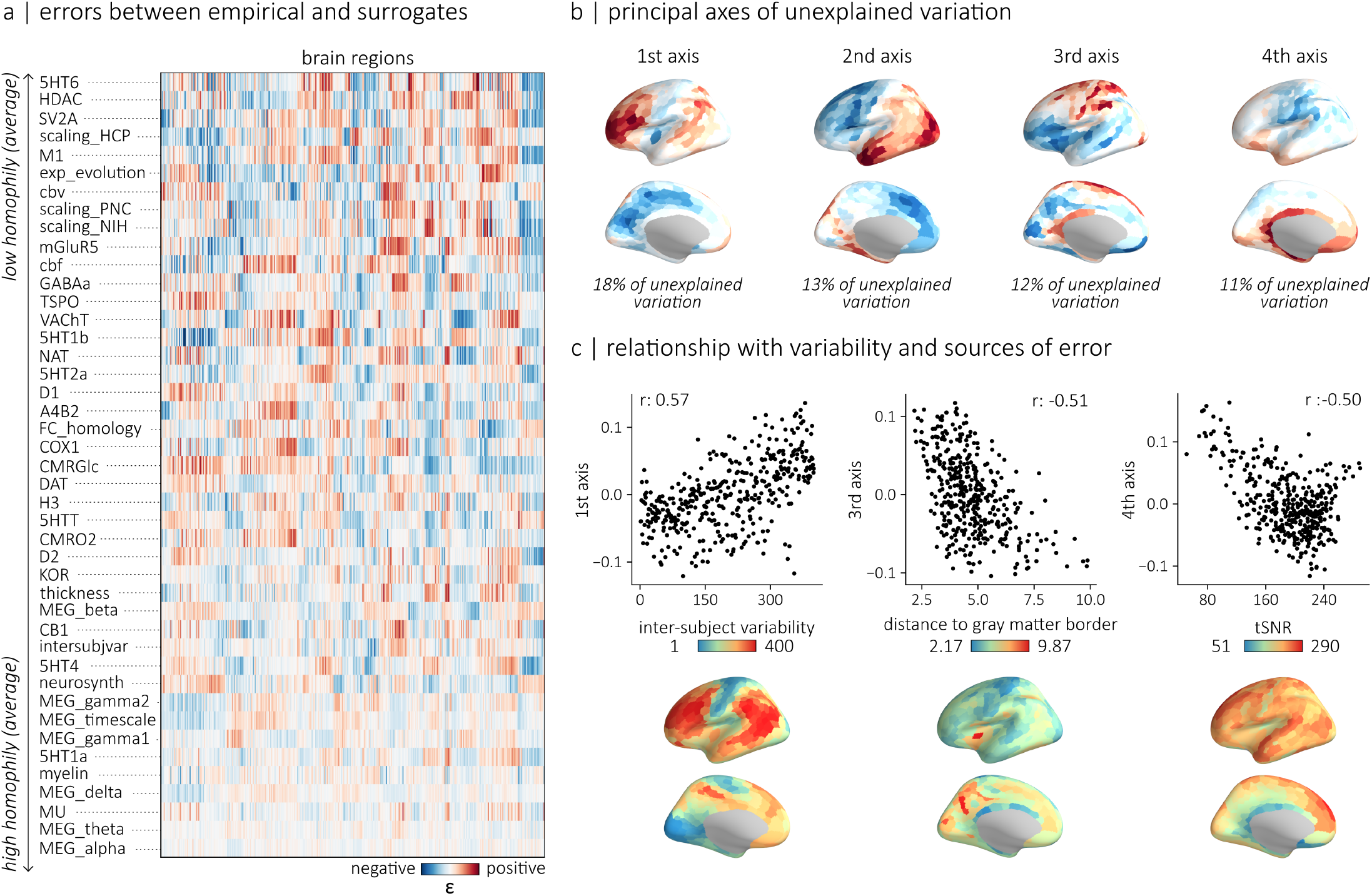
Extracting patterns of unexplained variation in brain maps. (a) Regional errors between empirical brain maps and their homophily-constrained surrogates, computed as the difference between ranked empirical values and the ranked mean of simulated maps. Brain maps are ordered by their average homophily across inter-regional modalities. (b) Principal axes of unexplained variation derived from the matrix of regional error, respectively explaining 18%, 13%, 12% and 11% of the variance. (c) Scatterplots capturing the relationship between axes of unexplained variation and potential sources of variability or error. Shown are the relationship between the 1^st^ axis of unexplained variation and intersubject variability in functional connectivity (left), the relationship between the 3^rd^ axis of unexplained variation and the parcel-averaged distance of each vertex to the gray matter border (middle) and the relationship between the 4^th^ axis of unexplained variation and the functional MRI temporal signal-to-noise ratio (tSNR) (right).

We find consistent error patterns, which suggests that they do not reflect map-specific noise but rather a common organizational structure across brain maps with low homophily. Indeed, we performed a principal component analysis on the matrix of regional errors and identified 4 axes of variance that capture, in total, 54% of the variance in the matrix (Fig. 4b). By definition, these error patterns are unrelated to the six modalities used to construct our simulated brain maps. They might however reflect additional meaningful modalities that were not considered in our analyses. To develop a better understanding of what these error patterns reflect, we calculated the component scores of each residual brain map (Fig. S4a). We also correlated the axes of variance with each brain map (Fig. S4b).

The 1^st^ axis of unexplained variation captures a gradient from medial brain regions to frontal and parietal regions on the lateral surface of the cortex. It is notably associated with the errors of maps capturing metabolic (e.g. CMRGlc and CMRO2) and functional (e.g. neurosynth, FC homology and cortical scaling) properties of brain regions. This error pattern might therefore be capturing variability along the functional axis of the cortex. Indeed, it is strongly associated with intersubject variability in functional connectivity (*r* = 0.57, *p*_S_ = 0.001; Fig. 4c, left) [29]. The 2^nd^ axis exhibits a rostro-caudal organization, from lateral temporal and occipital regions to medial prefrontal regions. The 3^rd^ axis is not as spatially smooth as the other axes and captures a gradient from ventral to dorsal cortical regions. Finally, the 4^th^ axis captures a gradient from medial to somatosensory regions.

Importantly, some of these error patterns may partly reflect acquisition- and preprocessing-related artifacts. The identification of such noise patterns is in itself interesting and desirable as they irrevocably influence downstream neuroimaging analyses. We find a strong correlation between the 3^rd^ axis and parcel-averaged distance of each vertex to the gray matter border (*r* = −0.51, *p*_S_ = 0.001; Fig. 4c, middle), suggesting that it may capture artifacts related to partial volume effects or registration inaccuracies. Similarly, the 4^th^ axis is strongly correlated with a map of fMRI-derived temporal signal-to-noise ratio (*r* = −0.50, *p*_S_ = 0.001; Fig. 4c, right), indicating that it may reflect regional variability in signal quality. Taken together, our results suggest that our error patterns may reflect both biologically meaningful and methodological sources of variation: some components may capture previously uncharacterized organizational principles whereas others may primarily reflect acquisition and preprocessing artifacts.

### Disentangling map-to-map associations using multimodal surrogates

The generative models introduced in this work can be used as surrogate maps to disentangle the effects of shared organizational structure when evaluating associations between cortical features. For example, they are well suited to examine the interplay between the wide array of cortical features that co-vary with the hierarchical axis of organization in the cortex [21]. We can take for instance the relationship between myelin and the principal axis of variance in functional connectivity [15](Fig. 5a, left). When controlling for spatial autocorrelation, we find a significant relationship between the two maps (*r* = −0.61, *p*_S_ = 0.001). However, myelin co-varies with a multitude of biological features, thus the significance of this correlation might arise as a result of general organizational rules driven by transcriptional, laminar and receptor similarity.

**Figure 5.**
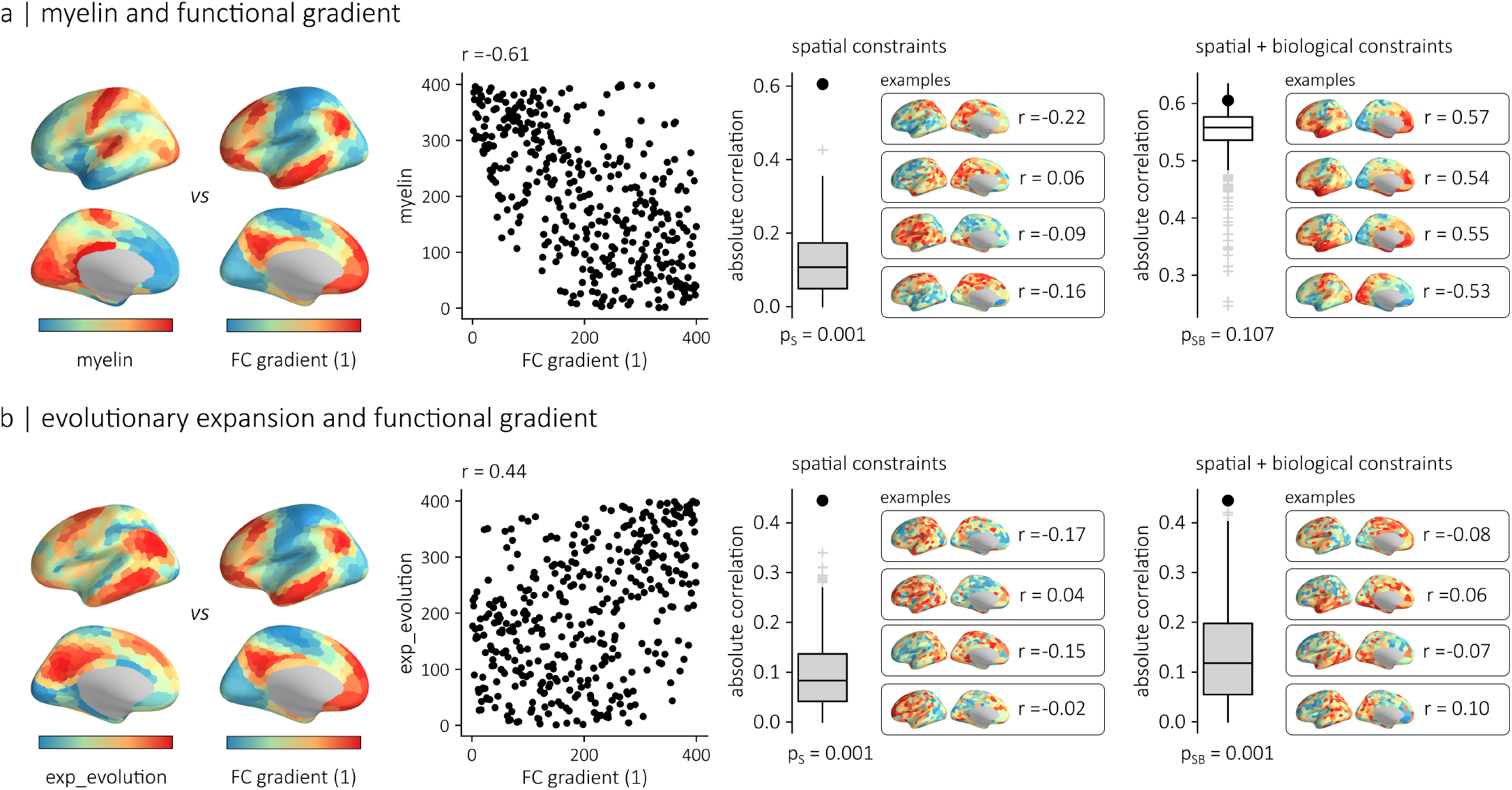
Using generative models to evaluate the interplay between biological constraints. (a) Relationship between cortical myelin and the principal functional connectivity (FC) gradient. Left: a scatterplot of ranked values shows a strong negative association between the two maps (*r* = *−*0.61). Middle: a null distribution of correlation coefficients obtained from surrogate maps preserving spatial autocorrelation (geodesic constraints) indicate that this association is significant (*p*_S_= 0.001). Right: when surrogate maps additionally preserve biological constraints (laminar, transcriptional, and receptor similarity), the association is no longer significant (*p*_SB_= 0.106). (b) Relationship between cortical evolutionary expansion and the principal FC gradient. Left: a scatterplot of ranked values shows a positive association between the two maps (*r* = 0.44). Middle: a null distribution of correlation coefficients obtained from surrogate maps preserving spatial autocorrelation (geodesic constraints) indicate that this association is significant (*p*_S_= 0.001). The association is still significant (*p*_SB_= 0.001) when the surrogate maps additionally preserve biological constraints (laminar, transcriptional, and receptor similarity).

To address this, we can generate surrogate myelin maps that preserve not only its spatial autocorrelation (i.e., Moran’s I with respect to spatial proximity), but also its Moran’s I with respect to each of the biological similarity matrices examined in this study (laminar, transcriptional and receptor similarity). These surrogate maps retain the structured relationship between myelin and these putative organizational forces, while removing its unique spatial configuration. By correlating these surrogates with the principal functional gradient, we can test whether the observed myelin–gradient association is explained by these shared organizational constraints. We find that the surrogate maps exhibit correlations with the principal functional gradient comparable to that of the empirical myelin map, resulting in a non-significant association between myelin and the principal functional gradient (*p*_SB_ = 0.106). In other words, these results show that this relationship can be accounted for by broader organizational principles potentially shared across multiple cortical features and may not reflect a “myelin-specific” association with the functional hierarchy.

We can similarly explore the relationship between cortical evolutionary expansion (macaque to human) and the principal axis of variance in functional connectivity (Fig. 5b). Indeed, previous findings suggest a strong relationship between the functional hierarchy, from unimodal sensory regions to multimodal association regions and the expansion of cortical regions [22, 30]. We reproduce this finding and identify a strong correlation between the two maps that is significant when controlling for spatial autocorrelation (*r* = 0.44, *p*_S_ = 0.001). We asked whether this relationship is explained by organizational constraints driven by transcriptional, laminar and receptor similarity between brain regions. We generated surrogate maps of evolutionary expansion preserving the spatial autocorrelation of the original map and its Moran’s I with respect to transcriptional, laminar and receptor similarity, and evaluated the significance of the correlation with respect to these surrogate maps. In this case, the correlation is still significant (*p*_SB_ = 0.001), which suggests that there is a unique link between these two maps.

## DISCUSSION

In this work we characterized the homophilic organization of a diverse set of brain maps and introduced a generative modeling framework that preserves homophily across biological, connectivity and geometric constraints. We showed that homophilic principles are sufficient to accurately reconstruct many empirical brain maps, with biological similarity and functional connectivity often contributing more than the brain’s geometry. We also identified consistent patterns of unexplained variation in maps with low homophily, revealing axes of cortical organization not captured by canonical interregional relationships. Finally, we showed that our generative model can disentangle the shared organization structure of brain maps and clarify whether associations between brain features reflect specific relationships or broader system-level constraints.

Proximal brain regions tend to exhibit similar biological properties. The principle of homophily is at the core of prominent neurodevelopmental theories. The chemoaffinity hypothesis indeed posits that local interactions between molecular and cellular signals influence the formation of synaptic connections [20]. Similarly, the protomap hypothesis [31] supposes that continuous gradients of signaling molecules [32, 33] influence neuronal fate in the embryonic ventricular zone while the protocortex hypothesis suggests that the fate of a cell is altered by its connectivity [34]. In other words, the idea that the organization of cortical features is shaped by inter-regional interactions imposed by molecular gradients and cortico-cortical connectivity is ubiquitous. Importantly, quantifying the autocorrelation of brain maps does not allow us to infer the biological mechanisms that give rise to the observed patterns in brain maps, which emerge from complex developmental and molecular processes [35, 36]. It does, however, offer a simple and convenient framework to quantify and articulate the interdependence between biological features.

Despite the widespread use of brain maps in neuroscience, attempts at modeling their whole-brain topographical organization remain uncommon. Existing frameworks have largely focused on modeling brain networks, such as functional connectivity [37–39] and, less commonly, structural connectivity [40– These efforts model derivatives of the original imaging data (in the form of MRI-, MEG-, PET-estimated matrices) rather than the original data itself (in the form of spatial maps). A small set of generative models for brain maps have been proposed but have mostly been used for building spatial autocorrelation-preserving null models [43, 44]. Although their primary use is to control for the smoothness of cortical data, these models implicitly capture a fundamental property of cortical organization and can provide valuable insight into the structure of brain maps. Here, we extend this framework by jointly modeling multiple sources of autocorrelation, integrating spatial, biological, and connectivity-derived constraints to better capture the principles governing brain map topography.

Specifically, we introduced a generative model that uses simulated annealing to generate surrogate brain maps that match empirical autocorrelation structures across multiple modalities. We can express Moran’s I as a weighted sum of squared eigenmode amplitudes [45, 46]. Thus, constraining the Moran’s I of a brain map is conceptually equivalent to constraining its coordinates in the eigenbases of weight matrices. This links our framework to recent eigenmode-based approaches for studying cortical organization and reconstructing brain maps (e.g. [47]). Here, rather than directly reconstructing empirical maps using eigenmodes, we impose a set of simple, interpretable homophilic rules and sample the broader space of admissible solutions compatible with the specified constraints. As such, we allow for degenerate solutions: distinct spatial configurations can satisfy the same constraints [48]. In this sense, the present generative model could potentially be extended to investigate subject-level variability in brain map organization and support the development of personalized “digital twin” brain models [49].

Surprisingly, although geodesic proximity alone provides substantial explanatory power for brain map organization, its contribution is markedly diminished once laminar, transcriptional, receptor, and connectivity-based similarities are taken into account. This suggests that the influence of the brain’s spatial embedding on cortical topography is to a large extent indirect: reflecting underlying biological constraints that are themselves spatially structured. In other words, the brain’s spatial layout does not arise arbitrarily from its geometry. Rather, its physical embedding imposes constraints on developmental processes, such as the minimization of wiring costs [50], axonal tension [51, 52], or the propagation of morphogens within a physical substrate [53, 54], which together shape the topographic organization of biological features on the cortical surface.

The capacity to model multiscale brain maps opens one underappreciated question: which biological features cannot be accurately explained using inter-regional similarity? We find that a subset of brain maps cannot be accurately simulated using the homophilic constraints considered here, suggesting that additional factors shape their organization. One possibility is that other biological constraints, such as neuropeptide signaling [55], vascular features [56], or metabolic [57] and molecular connectivity [58], play an important role. Another possibility is that inter-subject variability leads to discrepancies between empirical and modeled brain maps. The distribution of most biological features is indeed variable across individuals [59] and since we relied on group-averaged brain maps constructed from a limited set of subjects, some natural variability is expected. We notably demonstrate a strong association between the 1st axis of variance in the matrix of regional error and inter-subject variability in functional connectivity [29]. We also show that an alternative, non-biological, explanation is that some empirical brain maps contain consistent acquisition or preprocessing artifacts that cannot be reproduced by biologically-grounded models. This is particularly relevant for PET-derived maps, which are known to be affected by partial volume effects and other sources of spatial bias [60]. MRI-derived measures are not exempt from artificial noise either, with for example varying levels of fMRI temporal signal-to-noise ratio across the cortex [61]. In such cases, the lack of fit may reflect limitations associated with the acquisition modalities, rather than missing organizational principles, and can be informative in its own right.

The present modeling framework also sets the foundation for building biologically richer and more precise null models. Traditional null models in brain imaging typically preserve one dominant feature of cortical organization, for example spatial autocorrelation [43, 44, 62, 63], or nodal strength [64]. While effective for controlling specific sources of dependence, these approaches usually control for a single covariate. The present framework allows the generation of surrogate maps that preserve multiple features of organization. As such, it allows us to ask whether observed associations between brain maps reflect a specific relationship or emerge as a secondary consequence of a common organizational structure. The proposed framework is highly flexible and can be adapted to incorporate specific hypothesis-driven constraints. It therefore allows us to build more realistic benchmarks for hypothesis testing in whole-brain mapping [16, 65].

## METHODS

### Brain maps

We assembled 43 cortical brain maps from a library of reference maps available in the `neuromaps` toolbox [24]. Only reference maps covering both hemispheres and providing complete coverage across parcels were retained. Furthermore, when two reference maps capturing the same biological process were available, we prioritized those of higher quality. Namely, for serotonergic receptors, we prioritized maps from the high-resolution atlas of the human brain’s serotonin system [3] while for PET-derived metabolism maps, we prioritized those from [9]. When of similar quality, reference maps were averaged together, weighted by the sample size of each map. See Table 1 for a list of the 43 brain maps analyzed in this study. Each brain map was parcellated into 800 brain regions according to the Schaefer atlas [66]. Because the calculation of geodesic distance was restricted to a single hemisphere, and to focus on the intrinsic organization of brain maps independent of interhemispheric projections, parcellated data from the left and right hemispheres were combined and averaged to produce a symmetric, single-hemisphere representation of each brain map. Furthermore, to standardize distributions across modalities and reduce sensitivity to outliers, each brain map was converted to a rank-based representation. See Figure S1 for a visual representation of each map on the cortical surface.

### Inter-regional relationships

We constructed weight matrices capturing the strength of the relationships between brain regions across six distinct modalities. The data was parcellated into 800 brain regions according to the Schaefer functional atlas [66]. To focus on the intrinsic organization of brain maps independent of inter-hemispheric projections, and because the calculation of geodesic distance was restricted to a single hemisphere, only inter-regional relationships within the left-hemisphere were considered in our analyses.

#### Physical proximity (geodesic)

Geodesic distance, which corresponds to the shortest distance between a pair of points measured along the cortical surface, was first calculated between each vertex using Dijkstra’s algorithm on a graph representation of the pial cortical mesh of the fsaverage5 surface [96]. A parcel-to-parcel weight matrix was then constructed, where each entry (*i, j*) corresponds to the inverse of the average geodesic distance between every vertex in parcels *i* and *j*.

#### Connectivity (structural and functional)

Connectomes were generated using data from the Human Connectome Project S900 release [95]. Scans from N=327 unrelated participants (age 28.6 ± 3.73 years, 55% females) were used to reconstruct a consensus structural and functional connectome. Informed consent was obtained for all subjects (the protocol was approved by the Washington University Institutional Review Board as part of the HCP). The participants were scanned in the HCP’s custom Siemens 3T “Connectome Skyra” scanner, and the acquisition protocol included a high angular resolution imaging (HARDI) sequence and four resting state fMRI sessions. Briefly, the dMRI data was acquired with a spin-echo EPI sequence (TR=5,520 ms; TE=89.5 ms; FOV=210 × 180 mm^2^; voxel size=1.25 mm^3^; b-value=three different shells i.e., 1,000, 2,000, and 3,000 s/mm^2^; number of diffusion directions=270; and number of b0 images=18) and the resting-state fMRI data was acquired using a gradient-echo EPI sequence (TR=720 ms; TE=33.1 ms; FOV=208 × 180 mm^2^; voxel size=2 mm^3^; number of slices=72; and number of volumes=1,200). Additional information regarding the acquisition protocol is available at [95]. The data was then pre-processed according to the HCP minimal pre-processing pipelines [97].

Structural connectomes were reconstructed from the dMRI data using the MRtrix3 package [98]. Fiber orientation distributions were generated using a multi-shell multi-tissue constrained spherical deconvolution algorithm [99, 100]. The initial tractogram was generated with 40 million streamlines, with a maximum tract length of 250 and a fractional anisotropy cutoff of 0.06. Spherical-deconvolution informed filtering of tractograms (SIFT2) was used to reconstruct whole brain streamlines weighted by cross-section multipliers [101]. More information regarding the individual network reconstructions is available at [102].

A group consensus structural network was built such that the mean density and edge length distribution observed across individual participants was preserved [103]. The weights of the edges in the consensus networks correspond to the log-transform of the number of streamlines in the parcels, averaged across participants for whom these edges existed. A group-average functional connectivity matrix was constructed by concatenating the regional fMRI BOLD time series of all four resting-state sessions from all participants and computing the zero-lag Pearson correlation coefficient between each pair of brain regions.

#### Laminar similarity

Laminar similarity quantifies how similar pairs of cortical regions are in terms of their cellular distribution across the cortical laminae. We used cell-staining intensity profiles sampled across 50 equivolumetric surfaces (from the pial to the white matter surface) of the Big-Brain atlas, a high-resolution (20*µm*) histological reconstruction of a postmortem brain from a 65-year-old male [7, 104]. The data, registered to the fsaverage surface (164k vertices), was obtained from the BigBrainWrap toolbox [105] and parcellated into 800 cortical regions. The laminar similarity matrix was then calculated as the partial correlation between the intensity profiles of pairs of cortical regions, correcting for the mean intensity profile across regions. This matrix was further thresholded to retain only positive correlations.

#### Receptor similarity

Receptor similarity quantifies the similarity of cortical regions in terms of the density of their neurotransmitter receptor and transporters. Specifically, we parcellated 19 PET-derived brain maps of neurotransmitter receptors and transporters (see Table 1 for a list of the receptor and transporter used) and calculated, for pairs of brain region, the correlation (Pearson’s *r*) between the receptor profiles of each region. This receptor similarity matrix was then thresholded to retain only positive correlations.

#### Transcriptional similarity

The transcriptional similarity matrix quantifies the similarity of the transcription profiles of brain regions. We relied on the Allen Human Brain Atlas (AHBA; https://human.brain-map.org/)[1], which provides regional microarray expression data from six post-mortem brains (1 female, ages 24-57, 42.5 ± 13.38). The AHBA data was pre-processed and mapped to parcellated brain regions using the abagen toolbox (https://github.com/rmarkello/abagen) [2].

Pre-processing steps included updating the MNI coordinates of tissue samples to those generated via non-linear alignment to the ICBM152 template anatomy (https://github.com/chrisgorgo/alleninf). Also, microarray probe information was re-annotated for all genes using data provided by Arnatkeviciute and colleagues [106]. Then, probes were filtered by only retaining those that have a proportion of signal to noise ratio greater than 0.5. When multiple probes indexed the expression of the same gene, the one with the most consistent pattern of regional variation across donors was selected. Samples were then assigned to individual regions in the parcellations. If a sample was not found directly within a parcel, the nearest sample, up to a 2mm-distance, was selected. If no samples were found within 2 mm of the parcel, the sample closest to the centroid of the empty parcel across all donors was selected. To reduce the potential for misassignment, sample-to-region matching was constrained by hemisphere and gross structural divisions (i.e., cortex, subcortex/brainstem, and cerebellum, such that e.g., a sample in the left cortex could only be assigned to an atlas parcel in the left cortex). All tissue samples not assigned to a brain region in the provided atlas were discarded. Tissue sample expression scores were then normalized across genes using a scaled robust sigmoid function [107], and were rescaled to a unit interval. Expression scores were also normalized across tissue samples using the same procedure. Microarray samples belonging to the same regions were then aggregated by computing the mean expression across samples for individual parcels, for each donor. Regional expression profiles were then averaged across donors to obtain a single genes x brain regions matrix. From the 15,632 genes listed in this matrix, we only retained genes which are preferentially expressed in the brain and which showed a consistent pattern of regional variation across donors (i.e. differential stability greater than 0.5). Ultimately, a region x region matrix of transcriptional similarity was constructed by correlating the gene expression profiles of each pair of brain region. This matrix was thresholded to retain only positive correlations.

#### Moran’s I

We quantified homophily using Moran’s I, a global measure of spatial autocorrelation [25, 26]. Moran’s I can be defined as

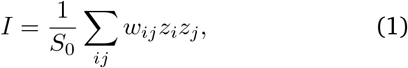

where *w*_*ij*_ corresponds to the weight quantifying the relationship between regions *i* and *j*. Larger weights indicate stronger relationships between pairs of regions. *S*_0_ corresponds to the sum of all weights *w*_*ij*_ and *z*_*i*_ corresponds to the standardized value of a brain map **u** in region *i*:

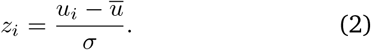

Moran’s I can be interpreted as a standardized crossproduct between a regional map and its neighborhood-averaged counterpart. Positive values indicate that regions tend to resemble their neighbors (positive autocorrelation; homophily), negative values indicate dissimilarity between neighboring regions (negative autocorrelation), and values near zero indicate no systematic relationship.

### Standardized Moran’s I

To enable comparison across modalities, Moran’s I was standardized using its expectation and variance under the null hypothesis that values were independently drawn from a normal distribution [26]. More specifically, the expected value of Moran’s I is:

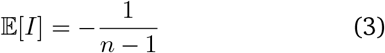

while the variance is defined as:

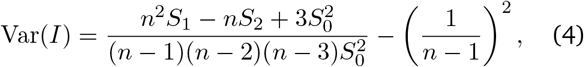

where

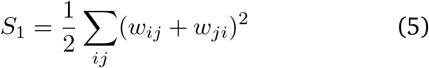

and

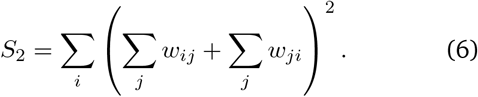

Ultimately, standardized Moran’s I is defined as:

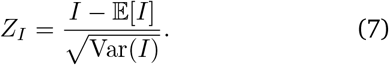

### Generative model

We introduce a generative model to simulate surrogate brain maps that preserve the homophilic (autocorrelation) structure of empirical maps across multiple weight matrices. The following procedure is described in terms of Moran’s I for simplicity. In this study, we use standardized Moran’s I (*Z*_*I*_) to account for differences in weight distributions and ensure that no weight matrix is favored in the generative process.

Given an empirical brain map **x** and 𝒲 = {**W**^(1)^, …, **W**^(*K*)^}, a set of *K* weight matrices, we start by computing a vector of Moran’s I values for map **x**:

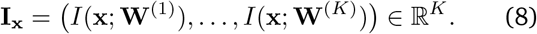

Our goal is to generate a surrogate brain map **y** such that

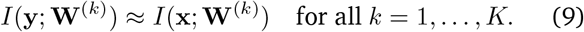

The algorithm starts with an initial random permutation of **x**:

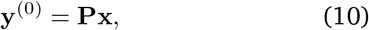

where **P** is a random permutation matrix. The algorithm then iteratively proposes swaps between pairs of indices *i* ≠ *j*, sampled uniformly at random, yielding random vectors **y**^*′*^. To accept or refuse the swap, we rely on a simulated annealing procedure. Namely, we define an energy function that quantifies the discrepancy between empirical and surrogate autocorrelation vectors. In this work, we define this energy function as the maximum absolute deviation (Chebyshev distance):

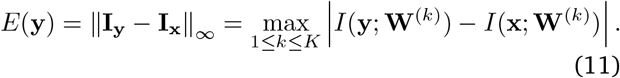

At iteration *t*, let *E*^(*t*)^ = *E* **y**^(*t*)^ and *E*^*′*^ = *E*(**y**^*′*^). The proposed swap is accepted with probability

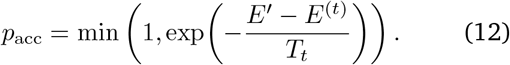

where *T*_*t*_ *>* 0 is the temperature parameter at iteration *t*. The temperature is gradually decreased according to a predefined cooling schedule. The algorithm is run until convergence, defined either by reaching a maximum number of iterations or by achieving *E*(**y**) below a specified tolerance. Ultimately, the resulting surrogate maps **y** preserve the empirical value distribution and, up to a specified tolerance, the empirical Moran’s I values across all weight matrices simultaneously.

### Efficiently updating Moran’s I

A naive implementation of the simulated annealing procedure described above would recompute Moran’s I from scratch after each proposed swap, which would be computationally prohibitive. Instead, we can exploit the fact that each swap is a simple permutation between two entries *i j* to efficiently compute Δ*I* = *I*(**y**^*′*^; **W**) − *I*(**y**; **W**).

In matrix form, Moran’s I can be written as:

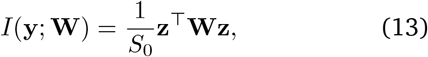

where *S*_0_ = Σ_*ij*_*w*_*ij*_ and where **z** is the standardized form of **y** (swapping entries in **y** is equivalent to swapping entries in **z**). *S*_0_ is constant across iterations. Thus, the only quantity that changes during the simulated annealing procedure is the quadratic form **z**^⊤^*W* **z**. Therefore, we can define the change induced by the swap between two entries *i* and *j* as:

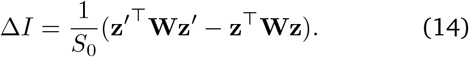

Importantly, the updated vector **z**^*′*^ differs from **z** only at positions *i* and *j* where 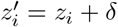, 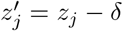, with δ = *z*_*j*_ − *z*_*i*_. Thus, by expanding the quadratic forms and canceling all terms unaffected by the swap, we have:

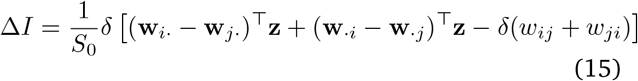

where **w**_*i*·_ and **w**_·*i*_ denote the i-th row and column of **W**, respectively. This equation assumes that *w*_*ii*_ = 0 for all *i*. Ultimately, evaluating Δ*I* only requires dot products between **z** and the row/column differences (**w**_*i*·_ − **w**_*j*·_) and (**w**_·*i*_ − **w** _· *j*_). This thus reduces the per-swap complexity from *O*(*n*^2^) to *O*(*n*).

### Reconstruction accuracy

To quantify how well the generative model reproduces the topography of empirical brain maps, we evaluated reconstruction accuracy as the similarity between empirical maps and their corresponding surrogate realizations. For each empirical brain map **x**, we generated 500 surrogate maps **y** using the generative procedure described above. Each surrogate preserves the distribution of empirical values (via permutation) and approximates its autocorrelation structure across all weight matrices.

Reconstruction accuracy was quantified as the absolute value of a Pearson correlation between the empirical map and each surrogate map. Our choice to use the absolute value reflects the fact that the generative model constraints the spatial structure of our maps (i.e. their autocorrelation) but does not enforce a specific orientation (i.e. sign).

### Model contributions

To quantify the relative importance of each interregional modality in shaping the organization of brain maps, we decomposed model performance into 4 distinct contribution metrics (Fig. S3a).

Let 𝒲= {**W**^(1)^, …, **W**^(*K*)^} denote the full set of *K* = 6 weight matrices. We evaluated the performance of our generative model under three configurations: (i) the *full* model, using all matrices in 𝒲 ; (ii) the *leave-one-out* models, in which a single matrix **W**^(*k*)^ is removed (i.e., 𝒲 \ **W**^(*k*)^); and (iii) the *marginal* models, in which only a single matrix **W**^(*k*)^ is used. For each configuration, we generated 500 surrogate maps and quantified reconstruction accuracy as described above.

The *leave-one-out* and *marginal* contributions of a modality *k* respectively correspond to the relative performance of the *leave-one-out* and *marginal* models compared to the *full* model:

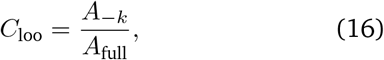

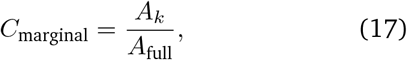

where *A*_full_ is the reconstruction accuracy of the full model, *A*_*−k*_ is the accuracy of the leave-one-out model excluding modality *k*, and *A*_*k*_ is the marginal accuracy obtained using modality *k* alone.

The *unique* contribution of a modality *k* captures the extent to which that modality provides information that is not redundant with the remaining modalities. We defined it as the relative decrease in model performance when **W**^(*k*)^ is removed from the full model:

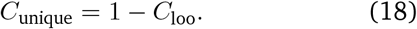

Larger values indicate that removing the modality substantially degrades performance, implying a greater unique contribution.

The *redundant* contribution quantifies the extent to which the explanatory power of a modality overlaps with that of other modalities. It is defined as the difference between the marginal and unique contributions:

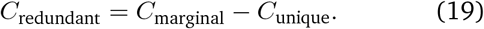

A large redundant contribution indicates that, although a modality can individually constrain brain map organization, much of its explanatory power is shared with other modalities in the full model.

### Low-dimensional morphospace

To visualize the relationships between brain maps and to contextualize the surrogate maps generated by our model, we constructed a low-dimensional morphospace using principal component analysis (PCA). PCA provides a global coordinate system in which distances between brain maps reflect their similarity structure, as evaluated using correlation-based metrics. This makes PCA preferable to nonlinear embedding techniques such as UMAP or t-SNE, which prioritize local neighborhood relationships and do not preserve global similarity or distance relationships.

We first assembled a data matrix **X** ∈ ℝ^*n×m*^, where *n* = 400 cortical regions (left hemisphere) and *m* = 43 empirical brain maps. Each column of **X** contained the rank-transformed values of a given brain map. Columns were mean-centered and scaled to unit variance across regions prior to decomposition. PCA was applied to **X**, and the first two principal components (PC1 and PC2), which together capture the greatest share of variance across maps, were used to define a two-dimensional morphospace. Each empirical map was projected onto these components to obtain its coordinates in this space. Surrogate maps were embedded in the same morphospace by projecting them onto the principal component axes derived from the empirical data. Surrogates were standardized in the same manner as the empirical maps before projection and to ensure that empirical and simulated maps have consistent orientations, we enforced a positive correlation between them. Namely, if the correlation was negative, the surrogate map was multiplied by −1.

### Axes of unexplained variance

To identify topographical patterns that cannot be captured by our generative model, we computed residual maps. For each empirical brain map **x**, we generated 500 simulated maps **y**. To ensure that empirical and simulated maps have consistent orientations, we enforced a positive correlation between them. Namely, if the correlation was negative, the surrogate map was multiplied by −1. We next calculated a residual map as the difference between the empirical map and the rank-transformed averaged surrogate map.

Residual maps were then assembled into a matrix **R** ∈ ℝ ^*n×K*^, where *n* is the number of cortical regions and *K* = 43 is the number of brain maps. We next performed principal component analysis (PCA) on **R** to identify dominant axes of shared residual variation across brain maps. Each component represents a spatial pattern of systematic deviation between empirical maps and model predictions. Component scores were obtained by projecting each residual map onto the resulting eigenvectors.

### Methodological covariates

To aid the interpretation of the axes of unexplained variance, we defined two metrics potentially associated with methodological sources of error: temporal signal-to-noise ratio (tSNR) and distance to the gray matter border.

#### Temporal signal-to-noise ratio

Temporal signal-to-noise ratio (tSNR) was estimated from the resting-state fMRI data used to construct the functional connectivity matrix. For each parcel of the Schaefer 800 atlas, tSNR was defined as the mean BOLD signal across time divided by the standard deviation of the corresponding time series. Regions with low tSNR are expected to be more sensitive to thermal and physiological noise sources.

#### Distance to the gray matter border

To estimate the susceptibility of cortical regions to partial volume effects and registration inaccuracies, we quantified the distance of cortical vertices to the gray matter border. A probabilistic gray matter mask for the MNI152NLin2009cAsym template was used to define the gray matter boundary. Gray matter vertices were defined as vertices with a gray matter probability greater than 0.05. For each vertex associated to parcels of the Schaefer 800 atlas (defined in MNI152NLin2009cAsym space, downloaded from TemplateFlow [108]), we computed the Euclidean distance to the nearest gray matter boundary voxel. Distances were then averaged across all vertices within each parcel to obtain a parcel-level measure of distance to the gray matter border. Larger values indicate regions located further from tissue-class boundaries and are therefore expected to be less susceptible to partial volume effects or registration inaccuracies.

### Significance testing using multimodal surrogates

To evaluate whether the association between two cortical maps can be explained by shared organizational constraints, we can compare an empirical correlation against a null distribution of correlations obtained from surrogate maps generated using our generative model. As an example, we evaluated the correlation between the principal functional gradient [15], and two brain maps capturing myelin content and cortical evolutionary expansion respectively. The cortical map capturing the principal functional gradient was obtained from neuromaps and pre-processed as described above (see *Brain maps* subsection).

For both myelin and cortical expansion, we generated 1000 surrogate brain maps only preserving Moran’s I with respect to geodesic proximity and 1000 surrogate brain maps preserving Moran’s I with respect to geodesic proximity, laminar similarity, transcriptional similarity, and receptor similarity. The resulting surrogates were then used to compute p-values controlling for spatial autocorrelation (*p*_*S*_) and p-values controlling for both spatial and biological autocorrelation (*p*_*SB*_).

Associations between cortical maps were quantified using Pearson’s correlation. Importantly, if a surrogate map **y** satisfies the autocorrelation constraints, then its sign-flipped counterpart (− **y**) also satisfies the same constraints. Consequently, the null distribution of correlation coefficients is symmetric around zero and bimodal. To account for this sign ambiguity, we quantified association strength using the absolute value of the correlation coefficient. Statistical significance was then assessed by comparing the association strength obtained with the empirical brain map to the association strengths obtained with surrogate maps. Two-tailed permutation p-values were calculated by quantifying the proportion of surrogate realizations whose absolute correlation magnitude deviated from the center of the null distribution at least as much as the empirical statistic.

## DATA AND CODE AVAILABILITY

The data and code used to conduct the analyses and generate the figures presented in this paper is available at https://github.com/netneurolab/bazinet_generative_model.

## ACKNOWLEDGMENTS

We thank Justine Y. Hansen, Eric. G. Ceballos, Yigu Zhou, Asa Farahani, Tahmineh Taheri and Moohebat Pourmajidian for their comments and suggestions on the manuscript. VB acknowledges support from the Natural Sciences and Engineering Research Council of Canada (NSERC), the Fonds de Recherche du Quebec - Nature et Technologie (FRQNT), the Healthy Brains for Healthy Lives initiative and Brain Canada. BM acknowledges support from the Natural Sciences and Engineering Research Council of Canada (NSERC), Canadian Institutes of Health Research (CIHR), Brain Canada Foundation Future Leaders Fund, the Canada Research Chairs Program, the Michael J. Fox Foundation, and the Healthy Brains for Healthy Lives initiative. The funders had no role in study design, data collection and analysis, decision to publish or preparation of the manuscript.

## AUTHOR CONTRIBUTIONS

Conceptualization: V.B., Z.Q.L., F.M., A.L. and B.M.; Methodology: V.B., Z.Q.L. and B.M.; Formal Analysis: V.B.; Writing - Original Draft: V.B. and B.M.; Writing - Review and Editing: V.B., Z.Q.L., F.M., A.L. and B.M.; Visualization: V.B. and Z.Q.L.; Supervision: B.M.

---

**Figure S1.**
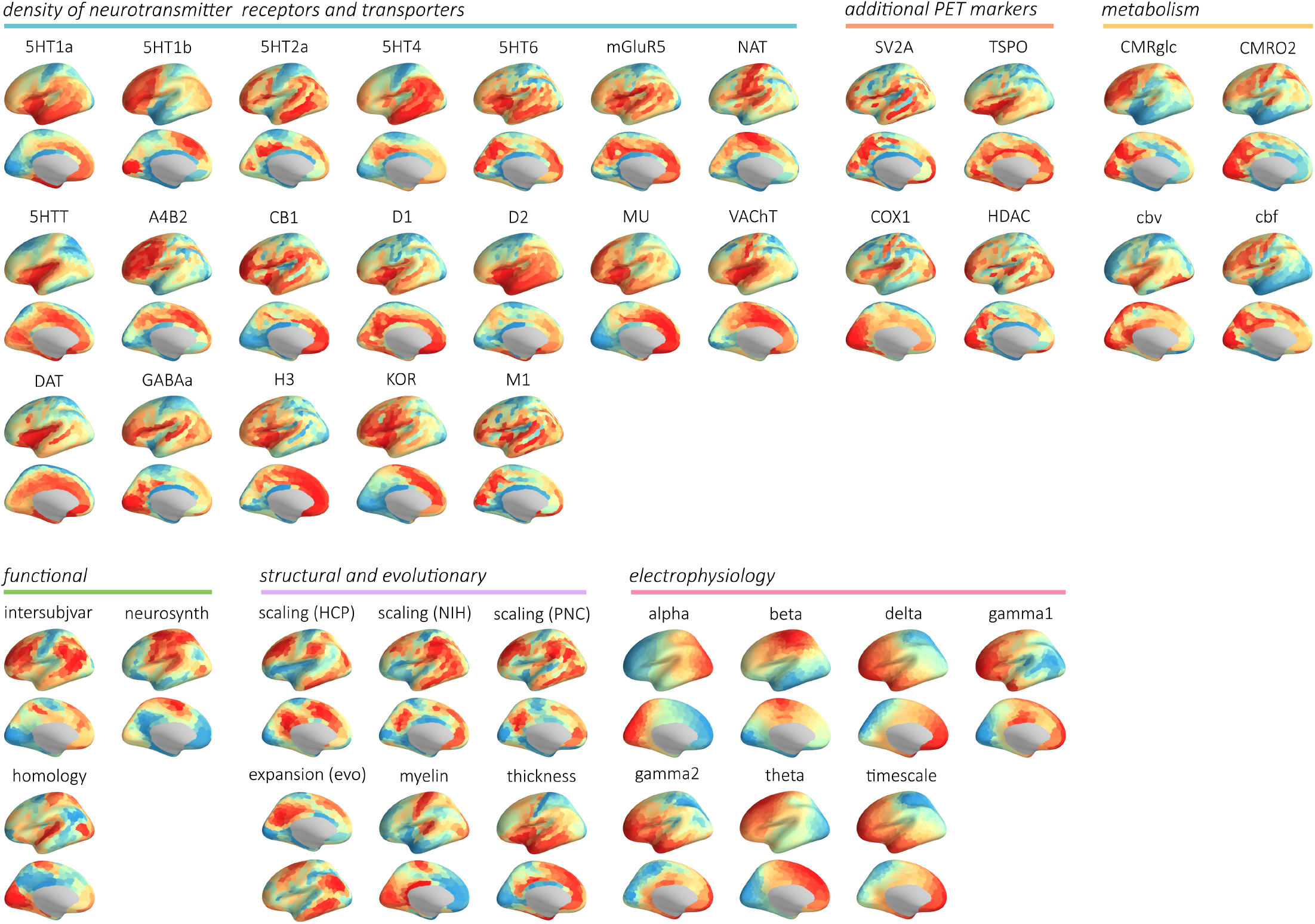
Cortical brain maps. Brain maps capturing the spatial organization of 43 features over the cortical surface. See Table 1 for a short description of each brain map and see *Methods* for more information on the methodology.

**Figure S2.**
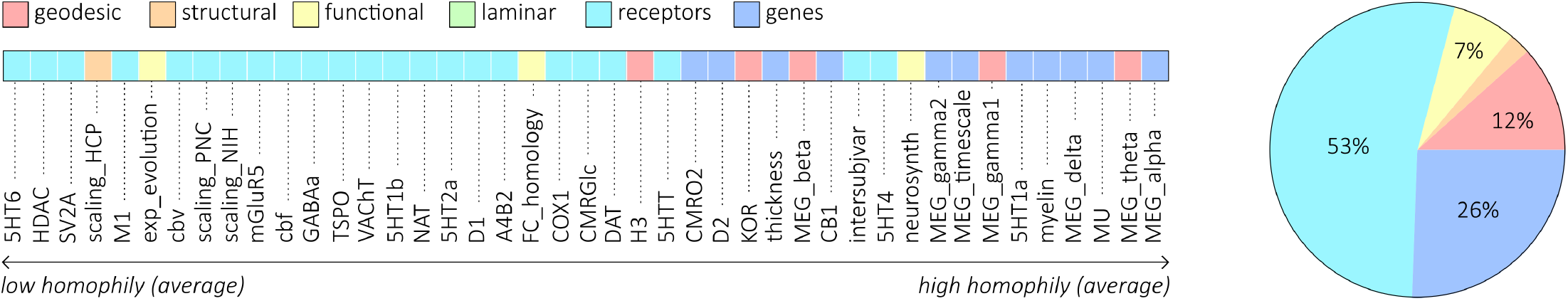
Modality associated with the highest homophily across brain maps. Left: brain maps are ordered according to their average autocorrelation across modalities, with colored labels indicating the modality associated with the largest standardized Moran’s I value. Right: proportion of brain maps for which each modality exhibited the largest standardized Moran’s I value. Most maps are more strongly autocorrelated with receptor similarity (53%) or gene transcription similarity (26%) than any other modality. Fewer maps are most strongly autocorrelated with geodesic proximity (12%), functional connectivity (7%), or structural connectivity (2%) than any other modality. No maps were more strongly autocorrelated with laminar similarity than any other modality.

**Figure S3.**
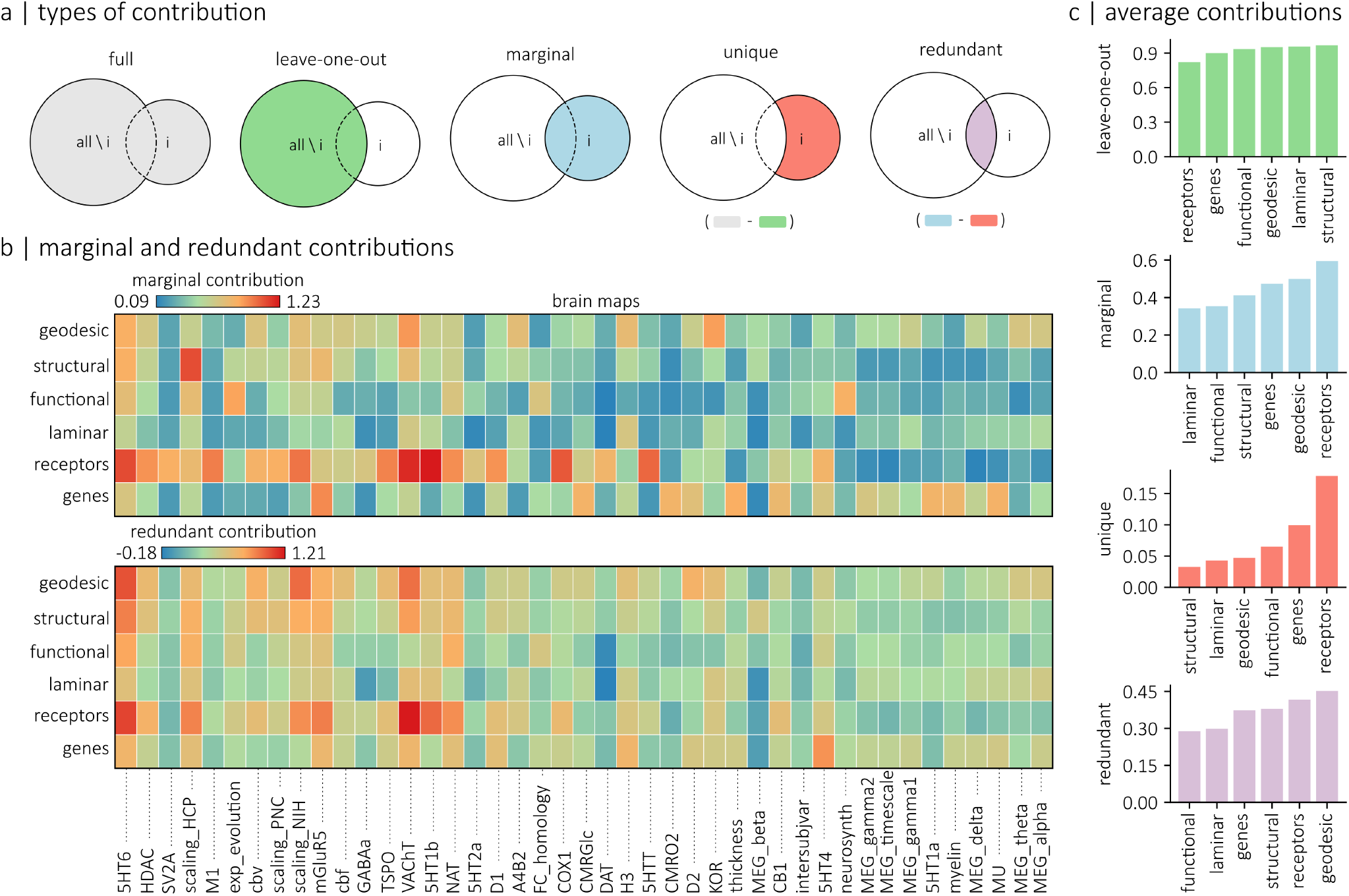
Contributions of inter-regional modalities to generative models of brain maps. (a) Schematic illustrating different types of contributions. The *full* contribution consists in the model’s performance when all modalities are included. The *leave-one-out* contribution captures the model’s performance when a specified modality is removed (*all \ i*). The *marginal* contribution reflects the model’s performance when a single modality is used in isolation (*i*). The *unique* contribution captures the difference in performance relative to the full model when a modality is excluded. Finally, the *redundant* contribution reflects the difference between the marginal and unique contributions of a modality. (b) top: marginal contributions of each inter-regional modality for each brain map. Bottom: redundant contributions of each inter-regional modality for each brain map. (c) Bar plot summarizing the average leave-one-out, marginal, unique, and redundant contribution of each modality across all brain maps.

**Figure S4.**
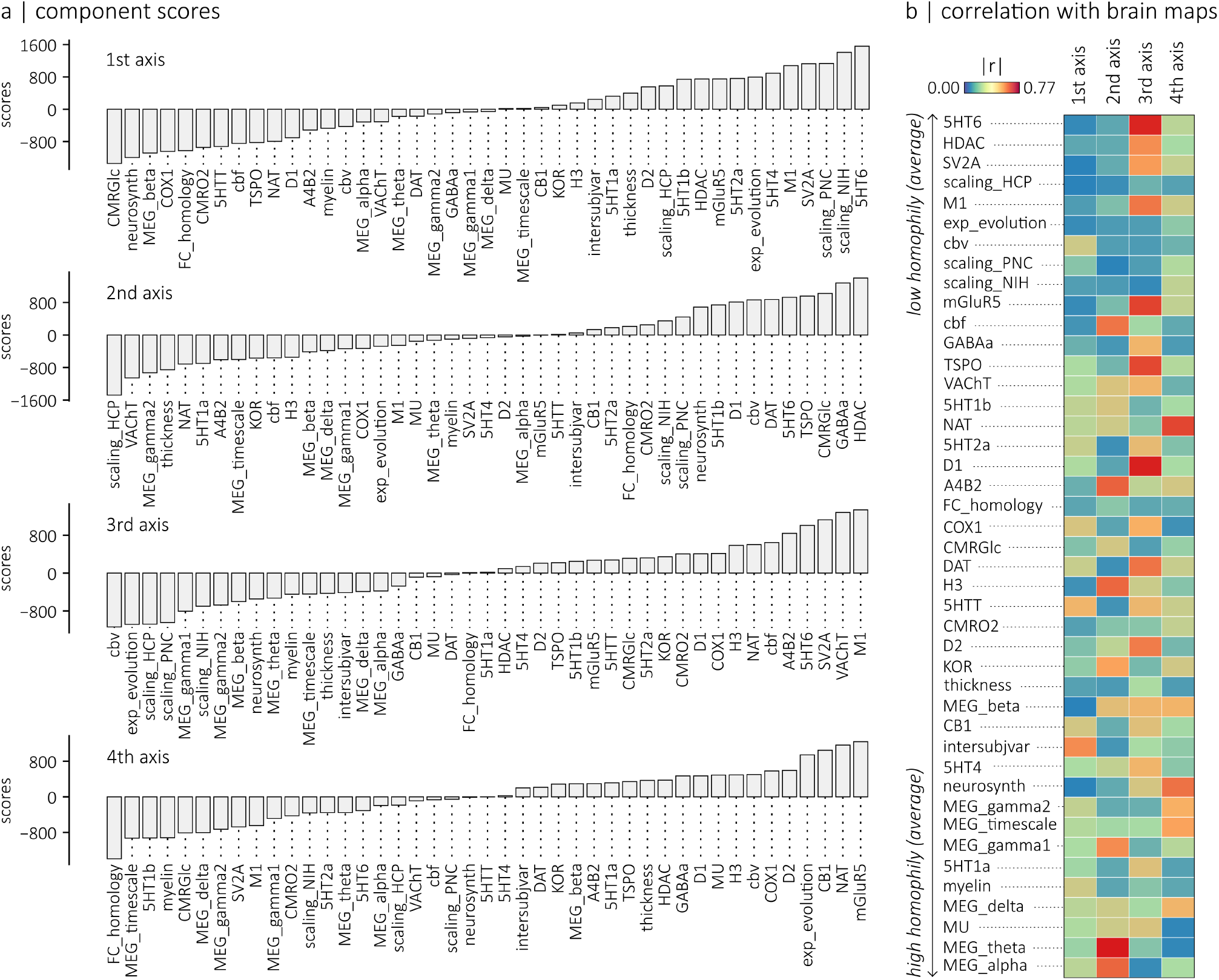
Relationship between axes of unexplained variation and empirical brain maps. (a) Component scores of each residual brain map, for each of the four principal axes of unexplained variation. (b) Heatmap showing the correlation between each empirical brain map and the principal components derived from the matrix of residual maps. Rows correspond to the first four axes of unexplained variation, and columns correspond to the 43 brain maps included in the analysis.

## References

[1] Hawrylycz, M. J. et al. An anatomically comprehensive atlas of the adult human brain transcriptome. Nature 489, 391–399 (2012).

[2] Markello, R. D. et al. Standardizing workflows in imaging transcriptomics with the abagen toolbox. eLife 10, e72129 (2021).

[3] Beliveau, V. et al. A high-resolution in vivo atlas of the human brain’s serotonin system. Journal of Neuroscience 37, 120–128 (2017).

[4] Hansen, J. Y. et al. Mapping neurotransmitter systems to the structural and functional organization of the human neocortex. Nature Neuroscience 25, 1569–1581 (2022).

[5] Glasser, M. F. & Van Essen, D. C. Mapping human cortical areas in vivo based on myelin content as revealed by t1-and t2-weighted mri. Journal of Neuroscience 31, 11597–11616 (2011).

[6] Alexander, D. C., Dyrby, T. B., Nilsson, M. & Zhang, H. Imaging brain microstructure with diffusion mri: practicality and applications. NMR in Biomedicine 32, e3841 (2019).

[7] Amunts, K. et al. Bigbrain: an ultrahigh-resolution 3d human brain model. Science 340, 1472–1475 (2013).

[8] Wagstyl, K. et al. Bigbrain 3d atlas of cortical layers: Cortical and laminar thickness gradients diverge in sensory and motor cortices. PLOS Biology 18, e3000678 (2020).

[9] Vaishnavi, S. N. et al. Regional aerobic glycolysis in the human brain. Proceedings of the National Academy of Sciences 107, 17757–17762 (2010).

[10] Castrillon, G. et al. An energy costly architecture of neuromodulators for human brain evolution and cognition. Science Advances 9, eadi7632 (2023).

[11] Frauscher, B. et al. Atlas of the normal intracranial electroencephalogram: neurophysiological awake activity in different cortical areas. Brain 141, 1130–1144 (2018).

[12] Gao, R., van den Brink, R. L., Pfeffer, T. & Voytek, B. Neuronal timescales are functionally dynamic and shaped by cortical microarchitecture. eLife 9, e61277 (2020).

[13] Mahjoory, K., Schoffelen, J.-M., Keitel, A. & Gross, J. The frequency gradient of human resting-state brain oscillations follows cortical hierarchies. eLife 9, e53715 (2020).

[14] Shafiei, G. et al. Neurophysiological signatures of cortical micro-architecture. Nature Communications 14, 6000 (2023).

[15] Margulies, D. S. et al. Situating the default-mode network along a principal gradient of macroscale cortical organization. Proceedings of the National Academy of Sciences 113, 12574–12579 (2016).

[16] Hansen, J. Y. & Misic, B. Integrating and interpreting brain maps. Trends in Neurosciences 48, 594–607 (2025).

[17] Hansen, J. Y. et al. Integrating multimodal and multi-scale connectivity blueprints of the human cerebral cortex in health and disease. PLOS Biology 21, e3002314 (2023).

[18] McPherson, M., Smith-Lovin, L. & Cook, J. M. Birds of a feather: Homophily in social networks. Annual Review of Sociology 27, 415–444 (2001).

[19] Wolpert, L. One hundred years of positional information. Trends in Genetics 12, 359–364 (1996).

[20] Sperry, R. W. Chemoaffinity in the orderly growth of nerve fiber patterns and connections. Proceedings of the National Academy of Sciences 50, 703–710 (1963).

[21] Huntenburg, J. M., Bazin, P.-L. & Margulies, D. S. Large-scale gradients in human cortical organization. Trends in Cognitive Sciences 22, 21–31 (2018).

[22] Sydnor, V. J. et al. Neurodevelopment of the association cortices: Patterns, mechanisms, and implications for psychopathology. Neuron 109, 2820–2846 (2021).

[23] Wang, X.-J. Macroscopic gradients of synaptic excitation and inhibition in the neocortex. Nature Reviews Neuroscience 21, 169–178 (2020).

[24] Markello, R. D. et al. Neuromaps: structural and functional interpretation of brain maps. Nature Methods 19, 1472–1479 (2022).

[25] Moran, P. A. Notes on continuous stochastic phenomena. Biometrika 37, 17–23 (1950).

[26] Cliff, A. & Ord, J. Spatial Processes: Models & Applications Pion, 1981).

[27] Hansen, J. Y. et al. Correspondence between gene expression and neurotransmitter receptor and transporter density in the human brain. NeuroImage 264, 119671 (2022).

[28] Burt, J. B. et al. Hierarchy of transcriptomic specialization across human cortex captured by structural neuroimaging topography. Nature Neuroscience 21, 1251–1259 (2018).

[29] Mueller, S. et al. Individual variability in functional connectivity architecture of the human brain. Neuron 77, 586–595 (2013).

[30] Xu, T. et al. Cross-species functional alignment reveals evolutionary hierarchy within the connectome. NeuroImage 223, 117346 (2020).

[31] Rakic, P. Specification of cerebral cortical areas. Science 241, 170–176 (1988).

[32] Rubenstein, J. L. et al. Genetic control of cortical regionalization and connectivity. Cerebral Cortex 9, 524–532 (1999).

[33] Bishop, K. M., Goudreau, G. & O’Leary, D. D. Regulation of area identity in the mammalian neocortex by emx2 and pax6. Science 288, 344–349 (2000).

[34] O’Leary, D. D. Do cortical areas emerge from a protocortex? Trends in Neurosciences 12, 400–406 (1989).

[35] O’Leary, D. D., Chou, S.-J. & Sahara, S. Area patterning of the mammalian cortex. Neuron 56, 252–269 (2007).

[36] Rakic, P., Ayoub, A. E., Breunig, J. J. & Dominguez, M. H. Decision by division: making cortical maps. Trends in Neurosciences 32, 291–301 (2009).

[37] Honey, C. J., Kötter, R., Breakspear, M. & Sporns, O. Network structure of cerebral cortex shapes functional connectivity on multiple time scales. Proceedings of the National Academy of Sciences 104, 10240–10245 (2007).

[38] Sanz Leon, P. et al. The virtual brain: a simulator of primate brain network dynamics. Frontiers in Neuroinformatics 7, 10 (2013).

[39] Breakspear, M. Dynamic models of large-scale brain activity. Nature Neuroscience 20, 340–352 (2017).

[40] Vértes, P. E. et al. Simple models of human brain functional networks. Proceedings of the National Academy of Sciences 109, 5868–5873 (2012).

[41] Betzel, R. F. et al. Generative models of the human connectome. NeuroImage 124, 1054–1064 (2016).

[42] Oldham, S. et al. Modeling spatial, developmental, physiological, and topological constraints on human brain connectivity. Science Advances 8, eabm6127 (2022).

[43] Burt, J. B., Helmer, M., Shinn, M., Anticevic, A. & Murray, J. D. Generative modeling of brain maps with spatial autocorrelation. NeuroImage 220, 117038 (2020).

[44] Koussis, N. C. et al. Generation of surrogate brain maps preserving spatial autocorrelation through random rotation of geometric eigenmodes. Imaging Neuroscience 3, IMAG–a (2025).

[45] de Jong, P., Sprenger, C. & van Veen, F. On extreme values of moran’s i and geary’s c. Geographical Analysis 16, 17–24 (1984).

[46] Dray, S., Legendre, P. & Peres-Neto, P. R. Spatial modelling: a comprehensive framework for principal coordinate analysis of neighbour matrices (pcnm). Ecological modelling 196, 483–493 (2006).

[47] Pang, J. C. et al. Geometric constraints on human brain function. Nature 618, 566–574 (2023).

[48] Edelman, G. M. & Gally, J. A. Degeneracy and complexity in biological systems. Proceedings of the National Academy of Sciences 98, 13763–13768 (2001).

[49] Katsoulakis, E. et al. Digital twins for health: a scoping review. NPJ digital medicine 7, 77 (2024).

[50] Bullmore, E. & Sporns, O. The economy of brain network organization. Nature Reviews Neuroscience 13, 336–349 (2012).

[51] Essen, D. C. v. A tension-based theory of morphogenesis and compact wiring in the central nervous system. Nature 385, 313–318 (1997).

[52] Hilgetag, C. C. & Barbas, H. Role of mechanical factors in the morphology of the primate cerebral cortex. PLOS Computational Biology 2, e22 (2006).

[53] Crick, F. Diffusion in embryogenesis. Nature 225, 420–422 (1970).

[54] Stapornwongkul, K. S. & Vincent, J.-P. Generation of extracellular morphogen gradients: the case for diffusion. Nature Reviews Genetics 22, 393–411 (2021).

[55] Ceballos, E. G. et al. Organization of neuropeptide systems in the human brain. Nature Neuroscience 29, 1212–1224 (2026).

[56] Farahani, A. et al. Mapping cerebral blood perfusion and its links to multi-scale brain organization across the human lifespan. PLOS Biology 23, e3003277 (2025).

[57] Yakushev, I., Drzezga, A. & Habeck, C. Metabolic connectivity: methods and applications. Current Opinion in Neurology 30, 677–685 (2017).

[58] Sala, A. et al. Brain connectomics: time for a molecular imaging perspective? Trends in Cognitive Sciences 27, 353–366 (2023).

[59] Gordon, E. M. et al. Precision functional mapping of individual human brains. Neuron 95, 791–807 (2017).

[60] Erlandsson, K., Buvat, I., Pretorius, P. H., Thomas, B. A. & Hutton, B. F. A review of partial volume correction techniques for emission tomography and their applications in neurology, cardiology and oncology. Physics in medicine and biology 57, R119–R159 (2012).

[61] Parrish, T. B., Gitelman, D. R., LaBar, K. S. & Mesulam, M.-M. Impact of signal-to-noise on functional mri. Magnetic Resonance in Medicine: An Official Journal of the International Society for Magnetic Resonance in Medicine 44, 925–932 (2000).

[62] Alexander-Bloch, A. F. et al. On testing for spatial correspondence between maps of human brain structure and function. NeuroImage 178, 540–551 (2018).

[63] Markello, R. D. & Misic, B. Comparing spatial null models for brain maps. NeuroImage 236, 118052 (2021).

[64] Milisav, F., Bazinet, V., Betzel, R. F. & Misic, B. A simulated annealing algorithm for randomizing weighted networks. Nature Computational Science 5, 48–64 (2025).

[65] Váša, F. & Mišić, B. Null models in network neuroscience. Nature Reviews Neuroscience 23, 493–504 (2022).

[66] Schaefer, A. et al. Local-global parcellation of the human cerebral cortex from intrinsic functional connectivity mri. Cerebral Cortex 28, 3095–3114 (2018).

[67] Naganawa, M. et al. First in human assessment of the novel m1 muscarinic acetylcholine receptor pet radiotracer 11c-lsn3172176. Journal of Nuclear Medicine (2020).

[68] Radhakrishnan, R. et al. Age-related change in 5-ht6 receptor availability in healthy male volunteers measured with 11c-gsk215083 pet. Journal of Nuclear Medicine 59, 1445–1450 (2018).

[69] DuBois, J. M. et al. Characterization of age/sex and the regional distribution of mglur5 availability in the healthy human brain measured by high-resolution [11c] abp688 pet. European Journal of Nuclear Medicine and Molecular Imaging 43, 152–162 (2016).

[70] Smart, K. et al. Sex differences in [11c] abp688 binding: a positron emission tomography study of mglu5 receptors. European Journal of Nuclear Medicine and Molecular Imaging 46, 1179–1183 (2019).

[71] Ding, Y.-S. et al. Pet imaging of the effects of age and cocaine on the norepinephrine transporter in the human brain using (s, s)-[11c] o-methylreboxetine and hrrt. Synapse 64, 30–38 (2010).

[72] Hesse, S. et al. Central noradrenaline transporter availability in highly obese, non-depressed individuals. European Journal of Nuclear Medicine and Molecular Imaging 44, 1056–1064 (2017).

[73] Normandin, M. D. et al. Imaging the cannabinoid cb1 receptor in humans with [11c] omar: assessment of kinetic analysis methods, test–retest reproducibility, and gender differences. Journal of Cerebral Blood Flow & Metabolism 35, 1313–1322 (2015).

[74] Laurikainen, H. et al. Sex difference in brain cb1 receptor availability in man. NeuroImage 184, 834–842 (2019).

[75] Kaller, S. et al. Test–retest measurements of dopamine d1-type receptors using simultaneous pet/mri imaging. European Journal of Nuclear Medicine and Molecular Imaging 44, 1025–1032 (2017).

[76] Aghourian, M. et al. Quantification of brain cholinergic denervation in alzheimer’s disease using pet imaging with [18f]-feobv. Molecular Psychiatry 22, 1531–1538 (2017).

[77] Bedard, M.-A. et al. Brain cholinergic alterations in idiopathic rem sleep behaviour disorder: a pet imaging study with 18f-feobv. Sleep Medicine 58, 35–41 (2019).

[78] Nørgaard, M. et al. A high-resolution in vivo atlas of the human brain’s benzodiazepine binding site of gabaa receptors. NeuroImage 232, 117878 (2021).

[79] Gallezot, J.-D. et al. Determination of receptor occupancy in the presence of mass dose:[11c] gsk189254 pet imaging of histamine h3 receptor occupancy by pf-03654746. Journal of Cerebral Blood Flow & Metabolism 37, 1095–1107 (2017).

[80] Hillmer, A. T. et al. Imaging of cerebral α4β2* nicotinic acetylcholine receptors with (-)-[18f] flubatine pet: Implementation of bolus plus constant infusion and sensitivity to acetylcholine in human brain. NeuroImage 141, 71–80 (2016).

[81] Sasaki, T. et al. Quantification of dopamine transporter in human brain using pet with 18f-fe-pe2i. Journal of Nuclear Medicine 53, 1065–1073 (2012).

[82] Dukart, J. et al. Cerebral blood flow predicts differential neurotransmitter activity. Scientific Reports 8, 1–11 (2018).

[83] Sandiego, C. M. et al. Reference region modeling approaches for amphetamine challenge studies with [11c] flb 457 and pet. Journal of Cerebral Blood Flow & Metabolism 35, 623–629 (2015).

[84] Jaworska, N. et al. Extra-striatal d2/3 receptor availability in youth at risk for addiction. Neuropsychopharmacology 45, 1498–1505 (2020).

[85] Turtonen, O. et al. Adult attachment system links with brain mu opioid receptor availability in vivo. Biological Psychiatry: Cognitive Neuroscience and Neuroimaging 6, 360–369 (2021).

[86] Kantonen, T. et al. Interindividual variability and lateralization of μ-opioid receptors in the human brain. NeuroImage 217, 116922 (2020).

[87] Vijay, A. et al. Pet imaging reveals lower kappa opioid receptor availability in alcoholics but no effect of age. Neuropsychopharmacology 43, 2539–2547 (2018).

[88] Finnema, S. J. et al. Kinetic evaluation and test–retest reproducibility of [11c] ucb-j, a novel radioligand for positron emission tomography imaging of synaptic vesicle glycoprotein 2a in humans. Journal of Cerebral Blood Flow & Metabolism 38, 2041–2052 (2018).

[89] Wey, H.-Y. et al. Insights into neuroepigenetics through human histone deacetylase pet imaging. Science translational medicine 8, 351ra106–351ra106 (2016).

[90] Kim, M.-J. et al. First-in-human evaluation of [11c] ps13, a novel pet radioligand, to quantify cyclooxygenase-1 in the brain. European Journal of Nuclear Medicine and Molecular Imaging 47, 3143–3151 (2020).

[91] Lois, C. et al. Neuroinflammation in huntington’s disease: New insights with 11c-pbr28 pet/mri. ACS Chemical Neuroscience 9, 2563–2571 (2018).

[92] Yarkoni, T., Poldrack, R. A., Nichols, T. E., Van Essen, D. C. & Wager, T. D. Large-scale automated synthesis of human functional neuroimaging data. Nature Methods 8, 665–670 (2011).

[93] Glasser, M. F. et al. A multi-modal parcellation of human cerebral cortex. Nature 536, 171–178 (2016).

[94] Reardon, P. et al. Normative brain size variation and brain shape diversity in humans. Science 360, 1222–1227 (2018).

[95] Van Essen, D. C. et al. The wu-minn human connectome project: an overview. NeuroImage 80, 62–79 (2013).

[96] Fischl, B., Sereno, M. I., Tootell, R. B. & Dale, A. M. High-resolution intersubject averaging and a coordinate system for the cortical surface. Human Brain Mapping 8, 272–284 (1999).

[97] Glasser, M. F. et al. The minimal preprocessing pipelines for the human connectome project. NeuroImage 80, 105–124 (2013).

[98] Tournier, J.-D. et al. Mrtrix3: A fast, flexible and open software framework for medical image processing and visualisation. NeuroImage 202, 116137 (2019).

[99] Jeurissen, B., Tournier, J.-D., Dhollander, T., Connelly, A. & Sijbers, J. Multi-tissue constrained spherical deconvolution for improved analysis of multi-shell diffusion mri data. NeuroImage 103, 411–426 (2014).

[100] Dhollander, T., Raffelt, D. & Connelly, A. Unsupervised 3-tissue response function estimation from single-shell or multi-shell diffusion mr data without a co-registered t1 image. In ISMRM Workshop on Breaking the Barriers of Diffusion MRI, vol. 5, 5 (2016).

[101] Smith, R. E., Tournier, J.-D., Calamante, F. & Connelly, A. Sift2: Enabling dense quantitative assessment of brain white matter connectivity using streamlines tractography. NeuroImage 119, 338–351 (2015).

[102] Park, B.-y. et al. Signal diffusion along connectome gradients and inter-hub routing differentially contribute to dynamic human brain function. NeuroImage 224, 117429 (2021).

[103] Betzel, R. F., Griffa, A., Hagmann, P. & Mišić, B. Distance-dependent consensus thresholds for generating group-representative structural brain networks. Network Neuroscience 3, 475–496 (2019).

[104] Paquola, C. et al. Microstructural and functional gradients are increasingly dissociated in transmodal cortices. PLOS Biology 17, e3000284 (2019).

[105] Paquola, C. et al. The bigbrainwarp toolbox for integration of bigbrain 3d histology with multimodal neuroimaging. eLife 10, e70119 (2021).

[106] Arnatkeviciute, A., Fulcher, B. D. & Fornito, A. A practical guide to linking brain-wide gene expression and neuroimaging data. NeuroImage 189, 353–367 (2019).

[107] Fulcher, B. D. & Fornito, A. A transcriptional signature of hub connectivity in the mouse connectome. Proceedings of the National Academy of Sciences 113, 1435–1440 (2016).

[108] Ciric, R. et al. Templateflow: Fair-sharing of multi-scale, multi-species brain models. Nature Methods 19, 1568–1571 (2022).

